# Optimizing Fed-Batch Processes with Dynamic Control Flux Balance Analysis

**DOI:** 10.1101/2024.06.11.598442

**Authors:** Mathias Gotsmy, Dafni Giannari, Radhakrishnan Mahadevan, Jürgen Zanghellini

**Affiliations:** Austrian Centre of Industrial Biotechnology, Austria; Department of Analytical Chemistry, University of Vienna, Austria; Department of Chemical Engineering and Applied Chemistry, University of Toronto, Canada

**Keywords:** bioprocess control, bi-level optimization, dynamic optimization, moving finite elements

## Abstract

Fed-batch processes are prevalent in biotechnological industries, but design of experiments often results in sub-optimal conditions due to incomplete solution space characterization. We employ a single-level dynamic control (DC) algorithm for dynamic flux balance analysis (dFBA), enhancing efficiency by reducing Karush-Kuhn-Tucker (KKT) condition constraints and adapting the algorithm for predicting optimal process length. In a growth-decoupled plasmid DNA production case study, we predict the optimal feeding profile and switching time between growth and production phase. Comparing our algorithm to its predecessor shows a speed-up of at least a factor of four. When the process length is part of the objective function the speed-up becomes considerably larger.

## 1. INTRODUCTION

Contrary, to the often cumbersome and expensive design of experiment studies, computer simulations of dFBA-based process models are a cheap and comparatively easy way to systematically sample the solution space for optimal fed-batch process designs [Gotsmy et al., 2023]. They can provide suggestions for setting control variables, for example, the feed rate [Espinel-Ríos et al., 2024] or the switch times of two-stage processes [Raj et al., 2020, Bauer and Klamt, 2024].

Recently, more sophisticated process optimization algorithms have been developed [Nakama and Jäschke, 2022, de Oliveira et al., 2023]. Most importantly, they employ KKT condition constraints and collocation techniques to translate the dFBA process models into an non-linear programming (NLP) problem which is solved with interior-point methods. Unfortunately, these algorithms come with a new set of challenges, especially their sensitivity to hyperparameters and improvable convergence.

Therefore, in this study, we adapt the dynamic control flux balance analysis (dcFBA) algorithm from de Oliveira et al. [2023]. Specifically, we reduce the number of KKT condition constraints and reformulate the finite element length constraints. These adaptations improve the general convergence speed as well as performance for (fed-batch) process length optimization.

## 2. THEORY

Here, we highlight the basic principles of our dcFBA implementation. For clarity, Table 1 lists the dimensions of vectors and matrices used throughout the section.

**Table 1.** Dimensions of vectors/matrices. *n*_R_ = *n*_𝔸𝔹ℂ𝔻_, number of reactions; *n*_M_, number of metabolites; *n*_FE_, number of finite elements.

| Variable | Shape | Variable | Shape |
| --- | --- | --- | --- |
| $\mathbf{S}$ | $(n_M, n_R)$ | $\alpha_{\text{lb}}, \mathbf{s}_{\text{lb}}$ | $(1, n_{\text{AB}})$ |
| $\mathbf{v}$ | $(n_R, 1)$ | $\alpha_{\text{ub}}, \mathbf{s}_{\text{ub}}$ | $(1, n_{\text{AC}})$ |
| $\mathbf{d}$ | $(1, n_R)$ | $\tau, \hat{\tau}$ | $(n_{\text{FE}}, 1)$ |
| $\lambda$ | $(1, n_M)$ | | |

### 2.1 Standard dynamic flux balance analysis

Fed-batch process simulations involve estimating state variables, such as biomass (*X*), substrate (e.g. glucose, *G*), product (*P* ), and volume (*V* ) from an initial state at a time *t*_0_ throughout the process length (*T* ). Differential equations for these state variables typically read

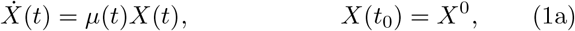

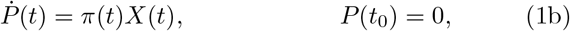

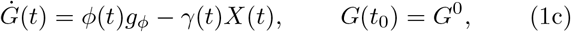

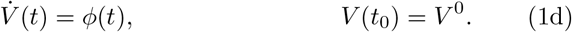

Here, 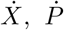, and *Ġ* represent the rates of change for biomass, product, and substrate, respectively. The parameters *μ, π*, and *γ* denote the specific growth rate, production rate, and substrate uptake rate, while *ϕ* represents the feed rate, and *g*_*ϕ*_ is the substrate concentration in the feed medium. For simplicity, we will assume *P* (*t*^0^) = *t*^0^ = 0. Additionally, for any time-dependent variable *x* at time *t*^*j*^, we adopt the shorthand notation *x*(*t*^*j*^) = *x*^*j*^.

In biotechnological production, engineers are typically interested in the optimization of key process performance metrics such as the final product titer

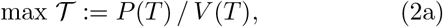

the average productivity

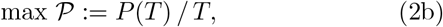

and the average product yield

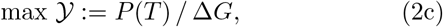

or combinations thereof [Zhuang et al., 2013], subject to the response of the cellular production host. Here, Δ*G* denotes the total substrate consumed.

Fed-batch production processes typically have several control variables (**u**) that can be set by the process engineer. Control variables, for example, comprise the feed rate [Dietzsch et al., 2011], the initial values of state variables [Gotsmy et al., 2023], or oxygen sparging and stirrer speed [Erian et al., 2018]. Thus, the DC problem asks: what values of **u** give an optimal process?

In dFBA, the host’s response to **u** is often approximated through parsimonious flux balance analysis (pFBA):

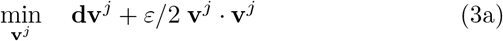

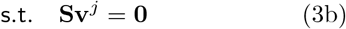

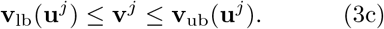

Here, **S** denotes the net stoichiometric matrix of the cell, **v**^*j*^ represents the cellular flux distribution, and **v**_lb_(**u**^*j*^) and **v**_ub_(**u**^*j*^) are the lower and upper bounds of the flux, respectively. The scaling parameter *ε* is typically assigned a small value [Ploch et al., 2020, Nakama and Jäschke, 2022, de Oliveira et al., 2023].

The host’s influence on the medium is evaluated in *n*_FE_ consecutive intervals *τ*^*j*^, and then aggregated across the entire process duration *T* :

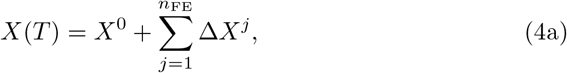

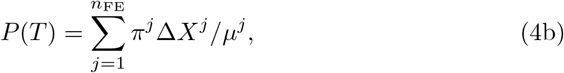

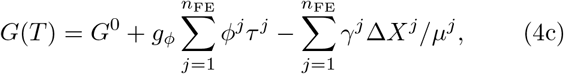

with

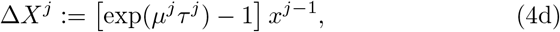

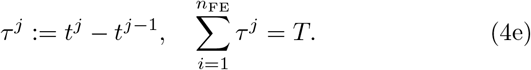

The changes in the amounts (Δ*X*^*j*^ and the summands in (4b) and (4c)) result from constant cellular growth with *μ*^*j*^ during the interval *τ*^*j*^. *μ*^*j*^, *π*^*j*^, and *γ*^*j*^ are a subset of the elements in the cellular flux vector **v**^*j*^.

Process (in)equality constraints depending on any system’s variable may be added to the optimization problem,

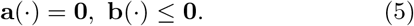

They typically comprise physical limitations (e.g., non-negativity of concentrations), technical limitations (e.g., lower and upper bound of the reactor volume or aeration rates), or biological limitations (e.g., maximal achievable biomass concentrations).

The system of equations (3), (4), and (5), combined with one of the objectives in (2), presents a bilevel optimization problem, which is generally challenging to solve [de Oliveira et al., 2023] even for state-of-the art algorithms like DFBAlab [Gomez et al., 2014].

### 2.2 Dynamic control FBA

dcFBA addresses this issue by transforming the bi-level optimization problem into a single non-linear optimization problem,

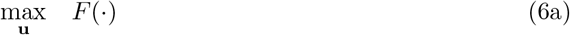

subject to

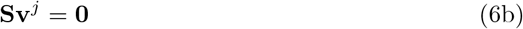

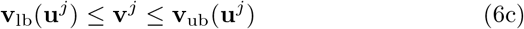

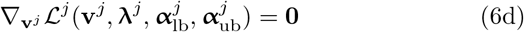

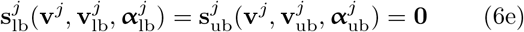

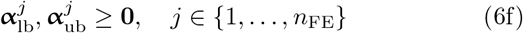

where the optimal solution 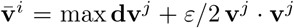 of the inner flux balance analysis (FBA) optimization in (3) satisfies the KKT conditions (6d)-(6f) [de Oliveira et al., 2023] with the Lagrangian

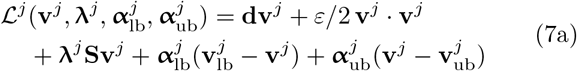

and

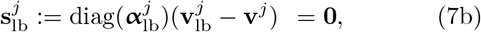

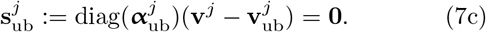

For numerical reasons, only KKT stationary conditions (6d) and KKT nonnegativity conditions (6f) are implemented as constraints. The KKT complementary slackness conditions (6e) are incorporated into the optimization function (6a) as a penalty term (𝒮) [de Oliveira et al., 2023]:

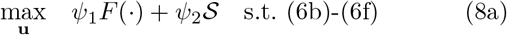

where

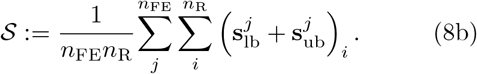

𝒮 converges to 0 from negative values, therefore, 𝒮 is maximized in the objective function. To achieve a balanced contribution of both terms, *F* and 𝒮, to the objective function, they are weighted by the hyperparameters *ψ*_1_ and *ψ*_2_.

Including 𝒮 as a penalty term has shown to overcome linear independence constraint qualification (LICQ) violations similar to the definition of the pFBA (Equation 3) [Baumrucker et al., 2008]. In accordance with previous studies, we assume that the LICQ holds for dcFBA [de Oliveira et al., 2023], however, a detailed analysis will be the focus of future work.

The single-level dcFBA optimization reformulation significantly improves optimization efficiency and thus renders more complex process optimization problems solvable [de Oliveira et al., 2023].

Our study is based on the work done by de Oliveira et al. [2023] and more detailed description of their dcFBA can be found there. Although their algorithm works well for short bioprocesses (i.e., *T* ≤ 10 h), its convergence becomes problematic for longer bioprocess, typically fed-batches, (i.e., *T* ≥ 30 h) if the temporal resolution (i.e. *n*_FE_*/T* ) is kept constant. Additionally, their implementation of finite element length bounds is impractical for process length optimization.

In the following sections, we focus on our adaptations to the algorithm. Henceforth, to increase the readability, we omit the index *j* from constraints when it is obvious.

### 2.3 Reducing KKT Condition Constraints

Previously, the KKT condition constraints of all fluxes of the metabolic model were included in the dcFBA algorithm [de Oliveira et al., 2023]. However, in metabolic modeling, for many fluxes no realistic constraints are known, usually indicated by lower bounds of -1000 and upper bounds of +1000 mmol g^−1^ h^−1^. The reactions of a metabolic model can be sorted into four classes, 𝔸 both bounds are known, 𝔹 only lower bounds are known, ℂ only upper bounds are known, and 𝔻 no bounds are known.

Here we reduce the number of process optimization constraints in our dcFBA algorithm by only including KKT condition constraints for reaction bounds that are known. Taking the partial derivative gives

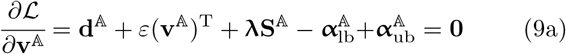

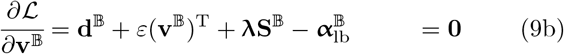

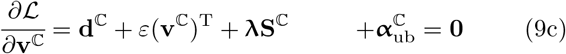

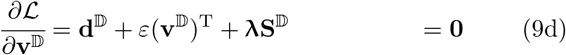

This method reduces the number of *α*-variables from 2*n*_R_ to 2*n*_𝔸_ + *n*_𝔹_ + *n*_ℂ_.

### 2.4 Rescaling Moving Finite Elements

In the previous approach, the lengths ***τ*** of finite element (FE) were not fixed but constrained by absolute lower and upper bounds. Yet, their sum added up to the total process length (*T* ),

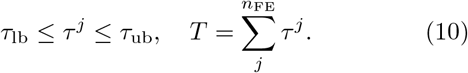

Setting upper and lower bounds on FE lengths serves two purposes: (1) enabling the optimization of total process length, and (2) managing sudden and strong rate changes during optimization [de Oliveira et al., 2023]. For biotechnological process optimization, both functionalities are important. Yet, relaxing the bounds on *τ*^*j*^ can lead to poor convergence, numerical issues, and even local infeasibility of the solver (see Section 4). Therefore, we limit the relative deviation of each FE from the average length *T/n*_FE_,

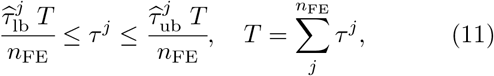

rather than the absolute length as in (10). Therefore, in our implementation, lower and upper bounds for *T* and 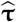 can be set separately for reasons (1) and (2), respectively. Our adaptation increases the number of unknown variables for optimization by one (i.e., the value for *T* ).

### 2.5 Adapted Dynamic control FBA

Finally, the derived constraints are integrated into our adapted dcFBA optimization problem,

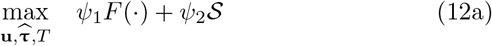

subject to FBA constraints

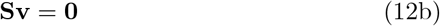

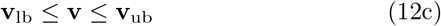

subject to KKT condition constraints

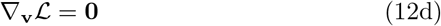

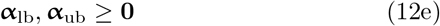

subject to moving FE constraints

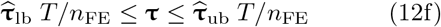

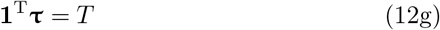

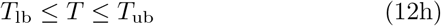

and subject to custom process (in)equality constraints

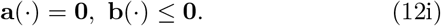

The KKT complementary slackness condition penalty term incorporated into the optimization function (12a) now reads:

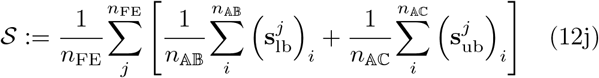

where *n*_FE_, *n*_𝔸𝔹_, and *n*_𝔸ℂ_ are used to normalize for the number of FEs and the number of lower and upper bound complementary slackness conditions, respectively.

## 3. METHODS

### 3.1 Plasmid DNA Production

In a prior study, we found that limiting sulfate in the reactor medium uncouples plasmid DNA (pDNA) production from biomass growth [Gotsmy et al., 2023]. Using dFBA, we designed and validated an optimal pDNA production process. Simulations were performed with the *E. coli* model *i*ML1515 [Monk et al., 2017] where pDNA synthesis was added expanding the model’s size to *n*_R_ = 2714, *n*_M_ = 1878.

To reduce the size of the genome-scale metabolic model (GSMM), we reran (classical) dFBA simulations for control and sulfate-limited processes from our prior study. At each evaluation time point, a lexicographic FBA was calculated followed by a pFBA. Reactions with no flux in any evaluation step were removed, resulting in a condensed GSMM with a size of *n*_R_ = 470 and *n*_M_ = 454.

Analysis of the experimental data [Gotsmy et al., 2023] showed that the pDNA synthesis rate (*π*) linearly decreased with process time and increased step-wise upon sulfate (*S*) limitation (Figure 1). For numerical reasons and differentiability, the step function was implemented as a tanh(·) with midpoint at *s* = *S/V* = 1 mmol L^−1^.

**Fig. 1.**
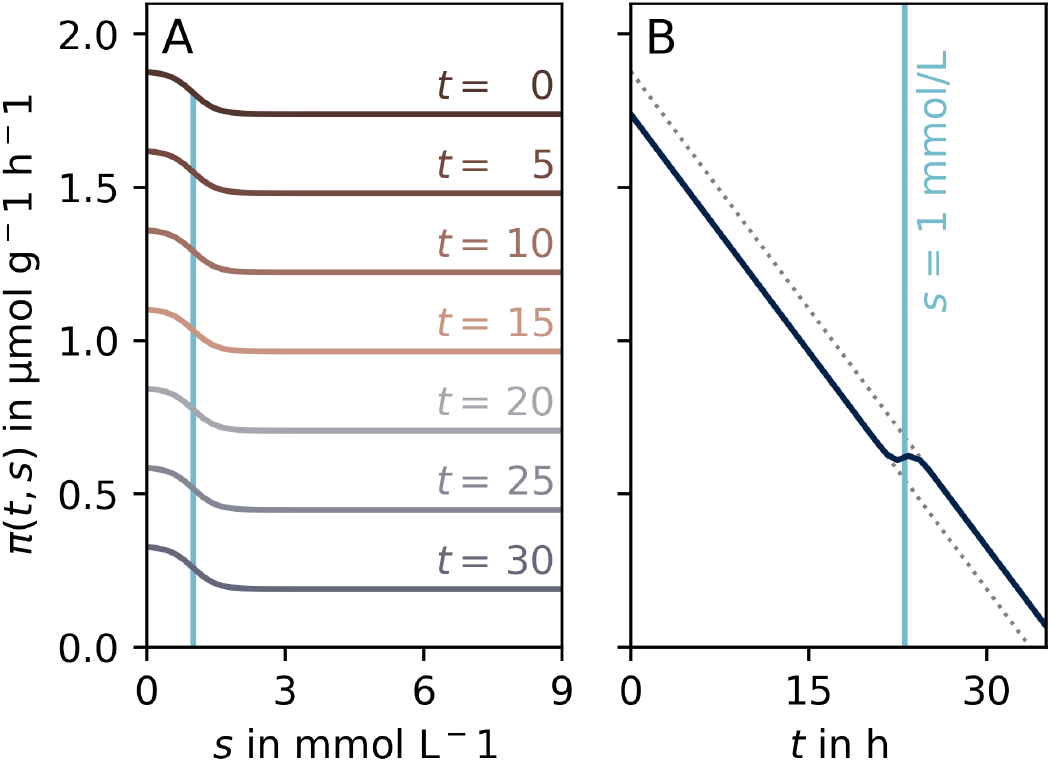
Production envelope of pDNA production. [Gotsmy et al., 2023]. *π*(*t, s*) is linearly dependent on fed-batch time (*t*) and increases by a step function during sulfate (*S*) starvation. **A** *π*(*t, s*) at different constant process times *t*. **B** *π*(*t, s*) of an example process where *s* = *S/V* = 1 mmol L^−1^ is reached after *t* = 23.1 h. The dotted lines indicate *π*(*t*) before and during *S* limitation.

### 3.2 Implementation of the Algorithm

The dcFBA algorithm was implemented following de Oliveira et al. [2023] using the Julia package JuMP [Lubin et al., 2023] and the nonlinear IPOPT solver [Wächter and Biegler, 2006]. Unless stated otherwise, all simulations used *n*_FE_ = 20 FEs and the IPOPT solver options were set as follows: `“tol” => 1e-4, “acceptable_iter” => 15, and “acceptable_tol” => 1e-2`.

The complete code for this study is available at https://github.com/Gotsmy/dcFBA.

## 4. RESULTS

In the following, we compare our adapted dcFBA implementation (referred to as `v2024`) with the previously published version [de Oliveira et al., 2023] (referred to as `v2023`). We conducted optimization for two pDNA production processes, aiming for maximal titer and maximal productivity using both algorithms.

Table 2 provides an overview of the algorithmic differences and optimization results for all four simulations. Columns *n*_𝔸_ to *n*_𝔹_ show the number of reactions per constraint class, as explained in the Methods section. In the `v2023` algorithm, all reactions are treated as fully constrained, despite their bounds being set to biologically unrealistic *±*1000 mmol g^−1^ h^−1^, placing them in class 𝔸. The number of dcFBA Lagrange multipliers (i.e, ***λ, α***_lb_, ***α***_ub_) per simulation is calculated as

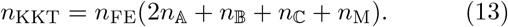

**Table 2.** Algorithmic Comparison of a previously published dcFBA algorithm ([de Oliveira et al., 2023], `v2023`) and our new method (`v2024`). We tested the optimization of titer (𝒯) and productivity (𝒫) in our pDNA production case study.

| Objective ($F$) | Algorithm | $n_A$ | $n_B$ | $n_C$ | $n_D$ | $n_{\text{KKT}}$ | Runtime [min] | $\mathcal{T}$ [g L<sup>-1</sup>] | $\mathcal{P}$ [g h<sup>-1</sup>] |
| --- | --- | --- | --- | --- | --- | --- | --- | --- | --- |
| $\mathcal{T}$ | v2024 | 1 | 290 | 0 | 179 | 14920 | 7.6 | 7.64 | 0.22 |
| $\mathcal{T}$ | v2023 | 470 | 0 | 0 | 0 | 27880 | 29.2 | 7.66 | 0.22 |
| $\mathcal{P}$ | v2024 | 1 | 290 | 0 | 179 | 14920 | 3.3 | 4.75 | 0.42 |
| $\mathcal{P}$ | v2023 | 470 | 0 | 0 | 0 | 27880 | 36.2 | 4.92 | 0.42 |

The corresponding column in Table 2 reveals that `v2024` reduces these multipliers by 54 % compared to `v2023` in this case study.

Optimizing the control problem involves optimizing each added multiplier during the simulation, contributing to computational costs. Notably, in the two optimization scenarios for productivtiy (𝒫) and titer (𝒯), both implementations converged to similar values. However, `v2024` outperforms `v2023`, being 11 and 4 times faster in the respective scenarios (see Table 2).

For the maximization of titer (𝒯), both `v2023` and `v2024` not only converged to the same values but also identified the same optimal process (Figure 2, light and dark blue line). Moreover, both `v2023` and `v2024` successfully predicted a sulfate-limited process for optimal productivity (𝒫) that shows identical characteristics to our previously experimentally implemented process [Gotsmy et al., 2023] (Figure 2, light and dark green line).

**Fig. 2.**
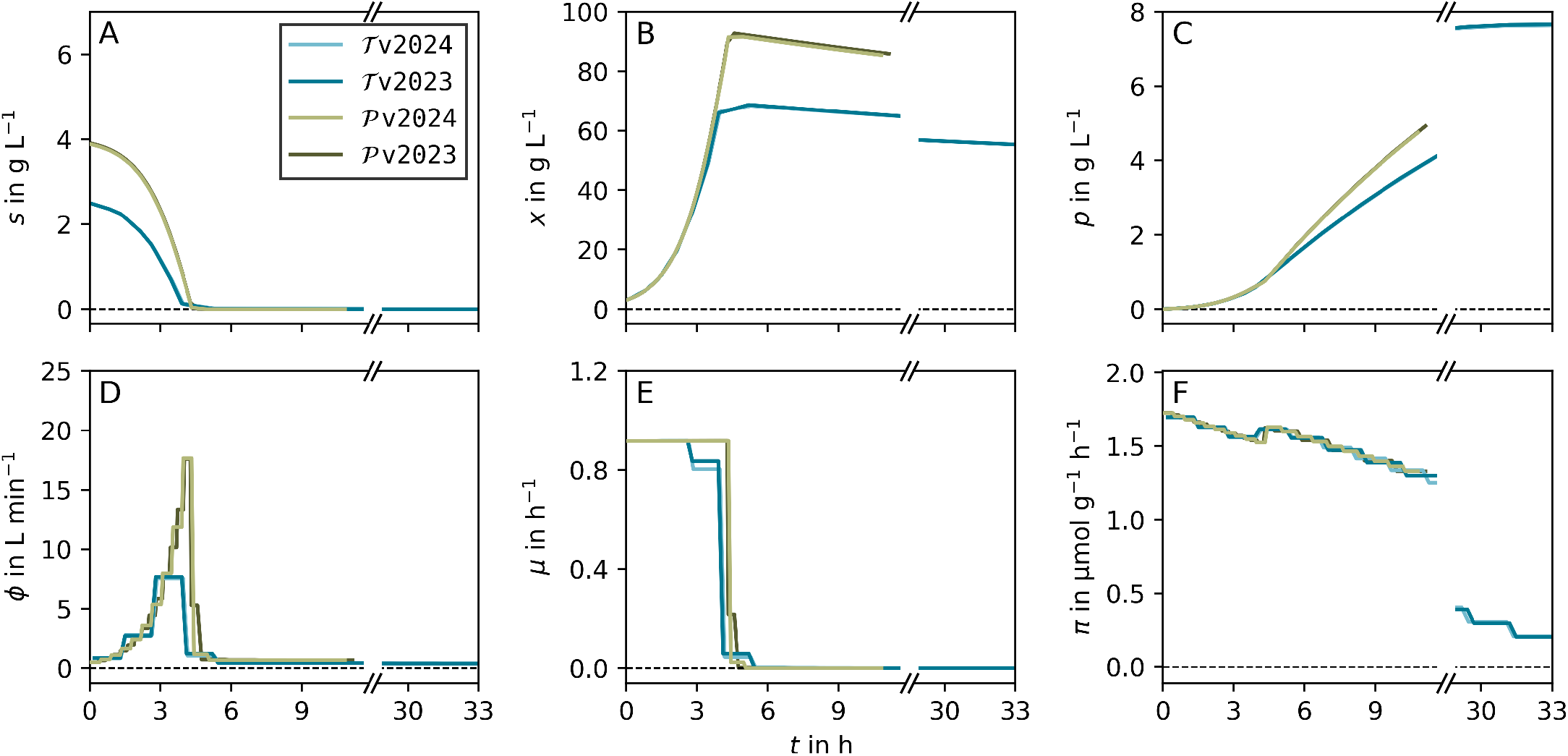
Comparison of the plasmid DNA (pDNA) production processes optimizing the titer (𝒯, blue) and the productivity (𝒫, green) by `v2023` (darker shade) and `v2024` (lighter shade), respectively. The top panels show integrated process state variables (sulfate *s*, biomass *x*, product *p*), and the bottom panels show rates (feed rate *ϕ*, growth rate *μ* and product synthesis rate *π*). The initial concentration of sulfate (panel A) and the feed rate (panel D) comprise the control variables (**u**) of the DC problem (Equation 12). For maximization of the titer both algorithms give the same result, however, there is a small difference for the maximization of productivity.

To analyze the difference between `v2024` and `v2023`, we investigated variations in the length distributions of FEs between both versions. Figure 3 compares FE lengths (*τ* ) for `v2023` (panel A) and `v2024` (panel B) when maximizing 𝒫.

**Fig. 3.**
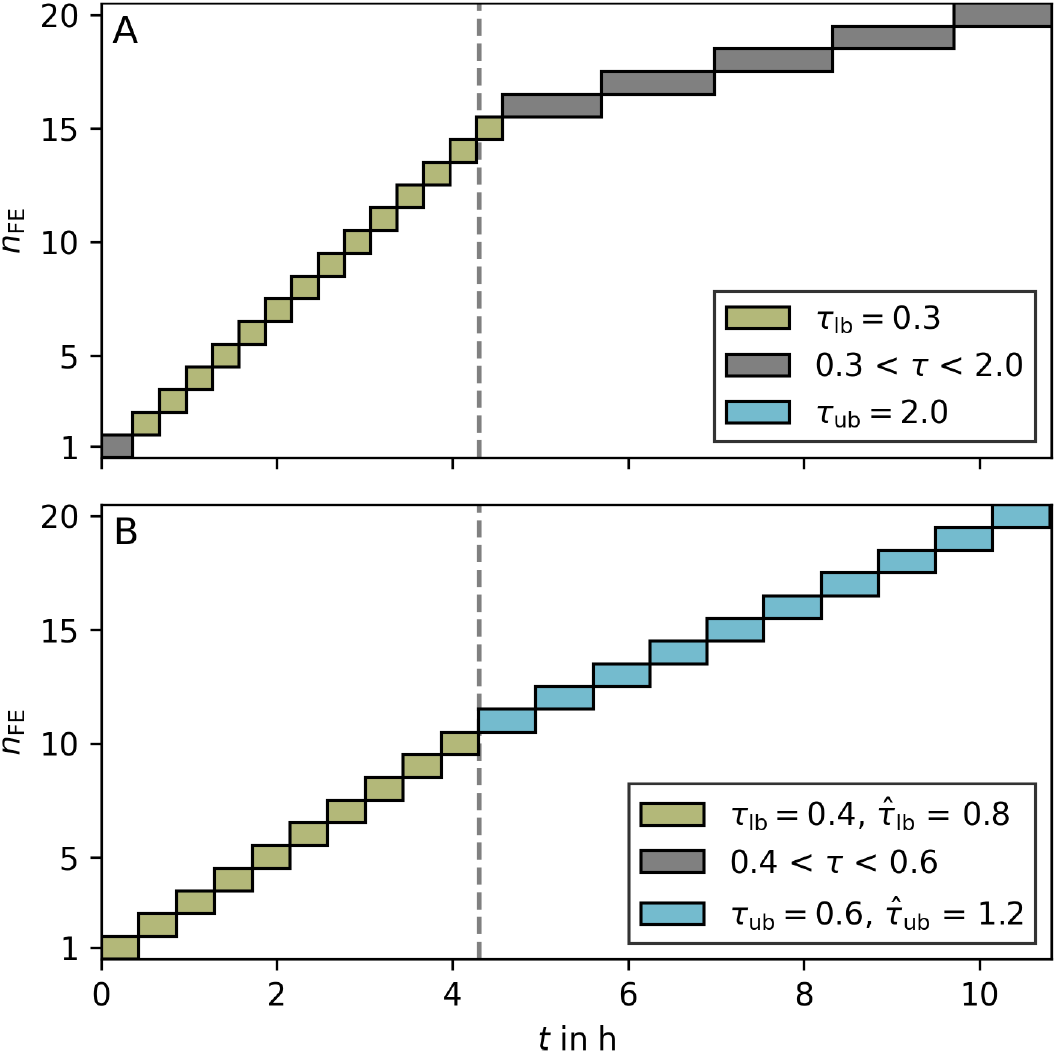
Comparison of FE lengths of 𝒫`v2023` (panel A) and 𝒫`v2024` (panel B). FEs colored green are equal to the lower length bound, FEs colored in blue are equal to the upper length bound, and grey FEs are in between. The grey dashed line indicates the switch from sulfate excess to sulfate limitation at *t* = 4.3 h.

Both versions exhibit a trend to shorter FE lengths initially, with FE becoming longer after growth cessation (*t* ≥ 4.3 h). However, in `v2023`, the average length of FE (1.1 h) after growth cessation is over 3.5 times larger than during the growth phase. This disparity leads to numerical and accuracy challenges, causing poor convergence of the IPOPT solver in `v2023`. In contrast, `v2024`, with its ability to independently set (tighter) bounds for *T* and *τ*^*j*^, exhibits smaller difference in FE lengths (Figure 3B), effectively avoiding numerical difficulties.

## 5. DISCUSSION

Designing optimal bioprocess is a grand challenge in industrial applications, often addressed by using dFBA. Optimizing bioprocesses with dFBA introduces computational complexities due to the bi-level nature of the problem. To mitigate this challenge, one strategy involves utilizing the KKT conditions, allowing for a transformation into a single-level but still non-linear optimization problem. Despite this simplification, the optimization remains computationally costly, with the attainment of a feasible solution heavily reliant on the algorithm’s hyperparameters.

Our results underscore these two critical considerations: (i) The necessity to identify and eliminate non-limiting or redundant constraints for accelerated performance. (ii) The importance of tightly controlling the algorithm’s hyperparameters to prevent numerical artifacts.

In standard GSMMs, reversible and irreversible reactions are commonly bounded within [-1000, 1000] and [0, 1000] mmol g^−1^ h^−1^, respectively, approximating the diffusion limit (private communication). These bounds are mostly active when the solution is otherwise unbounded, serving merely as indicators. This presents numerical challenges by inflating the number of constraints and computational costs, impacting both dFBA and FBA. While the importance of eliminating redundant constraints is recognized [Estinmgsih et al., 2019] and has been partially addressed [Nakama and Jäschke, 2022], the impact of default flux bounds has been overlooked. Our study demonstrates the effectiveness of removing these bounds; however, in rare cases, it may result in unbounded solutions. Utilizing formal methods [Estinmgsih et al., 2019] to identify redundant constraints mitigates this problem. Moreover, we believe that other approaches for dFBA for process optimization (e.g., [Scott et al., 2018]) or even the calculation of a standard FBA can profit from our proposed constraint reduction strategy.

While the inclusion of moving FEs in the DC problem increases its non-linearity, past cases have shown promising results [Gao et al., 2023]. However, without algorithmic incentives to minimize the discretization errors of the FEs, numerical errors may accumulate. Particularly when FE length bounds are broad due to process length optimization (Figure 3A). To mitigate this risk, setting bounds on relative FE lengths proves effective. This ensures a narrower distribution of FE lengths without overly constraining the total process length. Consequently, our algorithm is well-suited for optimizing bioprocesses where total process time (*T* ) is a part of the objective function.

Recently, an enhanced moving FE method which takes numeric errors into account has been developed [Gao et al., 2023], which will be part of future work.

## 6. CONCLUSION

Here we present an improved dynamic control flux balance analysis (dcFBA) algorithm. We showcase that strategic reformulation of the problem can decisively improve performance as well as convergence and accuracy compared to previous implementations. Overall, we hope that with careful initialization and selection of hyperprameters, this algorithm will offer a valuable tool for simulating and designing a wide range of bioprocesses.

## ACKNOWLEDGEMENTS

Authors acknowledge funding from the Canada Research Chairs Program to Radhakrishnan Mahadevan and the *Carbon–Cycle Economy Demonstration* (C-CED) project funded by the Austrian Climate and Energy Fund, carried out under the program “Vorzeigeregion Energie”, project number: 887638.

